# Coverage-preserving sparsification of overlap graphs for long-read assembly

**DOI:** 10.1101/2022.03.17.484715

**Authors:** Chirag Jain

## Abstract

Read-overlap-based graph data structures play a central role in computing *de novo* genome assembly using long reads. Many assembly tools use the string graph model [Myers, Bioinformatics 2005] to sparsify overlap graphs. Graph sparsification improves accuracy by removing spurious and redundant connections. However, a graph model must be coverage-preserving, i.e., it must ensure that each chromosome can be spelled as a walk in the graph, given sufficient sequencing coverage. This property becomes even more important for diploid genomes, polyploid genomes and metagenomes where there is a risk of losing haplotype-specific information.

We develop a novel theoretical framework under which the coverage-preserving properties of a graph model can be analysed. We first prove that de Bruijn graph and overlap graph models are guaranteed to be coverage-preserving. We also show that the standard string graph model lacks this guarantee. The latter result is consistent with the observation made in [Hui et al. ISIT’16] that removal of contained reads during string graph construction can lead to coverage gaps. To remedy this, we propose practical heuristics that are well-supported by our theoretical results to sparsify overlap graphs. In our experiments conducted by using simulated long reads from HG002 human diploid genome, we find that 50 coverage gaps are introduced on average by ignoring contained reads from nanopore datasets. We tested the proposed heuristics for deciding which contained reads should be retained to avoid the coverage gaps. The proposed method retains a small fraction of contained reads (1 – 2%) and closes majority of the coverage gaps.

## 1 Introduction

Long-read sequencing technologies have shown the greatest promise in computing high-quality haplotype-resolved genome assemblies [1,3,8,10,24,26,30,31,32]. Long-read accuracy has improved enormously over the last few years. For instance, PacBio HiFi sequencing technology yields consensus sequencing reads that are both long (averaging 10-25 kb) and highly accurate (averaging 99.8%) [29]. Similarly, modal raw read accuracy above 99% has been reported using reads from Oxford Nanopore Technology (ONT) sequencing [25]. Long and accurate reads are useful to sequence traditionally challenging diploid or polyploid genomes and facilitate haplotype-resolved *de novo* assembly [11,16,17,18]. The combination of PacBio and ONT reads has recently enabled the first complete gapless assembly of a human genome [22].

Genome assembly is the task of reconstructing the original sequence from a large number of reads. Read-overlap-based assembly algorithms work by constructing an *overlap graph* where each vertex corresponds to a read, and edges represent suffix-prefix overlaps between reads. Practical algorithms account for the following two challenges while computing genome assembly. First, it is usually possible to reconstruct different genomes using the same input set of reads, each of which is fully consistent with the data. Accordingly, problem formulations for genome assembly which seek a single genome reconstruction, e.g., by finding a Hamiltonian cycle in an overlap graph, or computing the shortest common superstring of input reads, are not used in practice [19]. Instead, assemblers compute contigs which are long, contiguous segments and unambiguously inferred to be part of the genome.

Second, the presence of genomic repeats and variable-length reads creates many spurious edges in the overlap graph. This challenge is often addressed by using graph sparsification heuristics that exclude vertices and edges which are likely to be redundant or false. The commonly used *string graph* model [21], for example, (i) removes reads that are *contained* as a substring of a longer read, (ii) ignores suffix-prefix overlaps that are shorter than the longest possible between a read pair, and (iii) removes transitive edges. More aggressive sparsification procedures also exist in the literature, for example, the *best overlap graph* approach proposed in [20] selects single best overlap per read end, where *best* is defined using some criterion. Inspite of the use of such graph sparsification techniques in most long-read assembly tools [3,4,6,7,12,13,23,28], theoretical understanding of these heuristics is fairly limited.

Assuming a genome is sequenced with sufficient coverage, then there must exist walks in an assembly graph that can spell the true sequences corresponding to each chromosome. An assembly graph which guarantees this property is said to be *coverage-preserving* (formally defined in the next section). Note that this property is not intended to check if the true sequences can be unambiguously reconstructed [2,27], but to check whether their information is preserved in the graph. In this paper, we make the following contributions:

1. We formulate the coverage-preserving property of genome assembly graph models, and perform a rigorous evaluation in three commonly-used models: (a) de Bruijn graphs, (b) overlap graphs and (c) string graphs. We prove that de Bruijn graphs and overlap graphs are guaranteed to be coverage preserving, but string graphs are not.
2. Graph sparsification is a critical step during genome assembly to prune the overlap graph, as it helps to compute longer contigs. We develop theoretical results to compute a sparse overlap graph while preserving the coverage-preserving property.
3. We extend the proposed theory into practical heuristics and a prototype software ContainX (github.com/ at-cg/ContainX). The proposed heuristics are useful to identify redundant contained reads, i.e., the contained reads which can be excluded from overlap graph without violating the coverage-preserving property. We empirically show that the proposed heuristics can help improve assembly quality.
4. We conducted experiments by using error-free long read datasets simulated from diploid HG002 human genome assembly while matching read length distributions to real PacBio HiFi and ONT data. We show that 1 and 50 coverage gaps are introduced on average by removing all contained reads in HiFi and ONT datasets respectively. ContainX algorithm excludes 98 – 99% contained reads from the graph and leaves ≤ 5 coverage gaps. This result compares favorably to the performance of other known solutions for this problem.

## 2 Preliminaries

### Notation on strings

For a linear string *x* = *a*_1_ … *a*_*n*_ over alphabet *Σ* = {A,C,G,T}, |*x*| = *n* is the length of *x*, *x*[*i*] = *a*_*i*_ is the *i*^*th*^ symbol of *X*, and *x*[*i* : *j*] = *a*_*i*_ … *a*_*j*_ is the substring from position *i* to position *j*. Let *x*^*i*^ denote string *x* concatenated with itself *i* times. Define *pre*_*i*_(*x*) to be the first *i* characters of string *x*. String *x* is said to have a suffix-prefix overlap of length *l* with string *y* when a proper suffix of length *l* in *x* equals a proper prefix of *y*. Recall that a proper prefix or a proper suffix of a string cannot be equal to the string itself and is non-empty.

### Circular genome model

We will use a circular genome model to avoid edge-effects associated with sequencing coverage. A circular string can be viewed as a traditional linear string which has the left- and the right-most symbols wrapped around and glued together. Circular string *z* = ⟨*a*_1_ … *a*_*n*_⟩ has length |*z*| = *n*. Substring *z*[*i* : *j*], where *i* ∈ [1, *n*], *j* ≥ *i*, equals the finite substring of the linear infinite string (*a*_1_ … *a*_*n*_)^∞^ from position *i* to position *j*. Note that offset *j* in *z* [*i* : *j*] can be greater than |*z*| because *z* is a circular string. Further, substring *z*[*i* : *j*] is said to be *repetitive* iff there exists *z* [*i*′ : *j*′] = *z* [*i* : *j*], *i*′ ≠ *i*. Suppose the true unknown genome is a set of circular strings, each representing a chromosome sequence. Let *ϕ* be a known upper bound on the maximum length of a chromosome in the genome. A sequence *read* sampled from the genome is a substring of one of the circular strings in the genome. An indexed multiset of the reads is represented using symbol 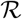. Read 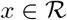 is labeled as a *contained* read if there exists a read 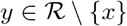 such that *x* is a substring of *y*. If *y* is also contained in *x*, then *x* and *y* are identical. In this case, break the tie by assuming that the read with lower index in 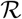 is contained within the read with higher index. The reads which contain read *x* are referred to as *parent* reads of *x*. The proposed framework assumes that there are no sequencing errors or reads have been error-corrected. In our theoretical results, we will also assume that DNA is a single-stranded molecule.

### Assumption on sequencing coverage

Depth of sequencing must be sufficient enough for computing *de novo* genome assembly. We wish to ensure that reads “cover” the genome, and the length of suffix-prefix overlap between “consecutive” reads is above a certain threshold. One way to achieve this is following. We say that there is *sufficient coverage* over circular string *z* if there exist parameters *l*_1_, *l*_2_ ∈ ℕ such that *l*_1_ > *l*_2_ and all intervals of length *l*_1_ – *l*_2_ in *z* include the start of at least one substring of length ≥ *l*_1_ that matches a read ∈ 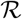. See Figure 1 for an example. Let 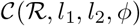 be the set of all *candidate* circular strings of length ≤ *ϕ* which satisfy the above coverage assumption using a given read set 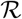 and coverage parameters (*l*_1_, *l*_2_). The genome is a subset of 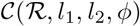.

**Fig. 1.**
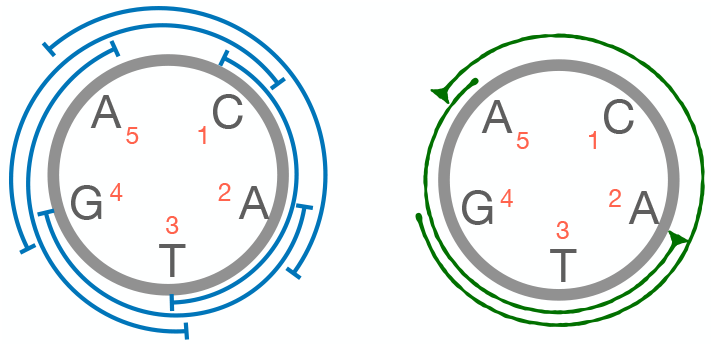
Suppose *l*_1_ = 4 and *l*_2_ = 1. A circular string *z* = ⟨CATGA⟩ has five intervals of length *l*_1_ – *l*_2_ = 3 as shown in blue. Two reads ACATG and ATGA, which match substrings *z*[5 : 9] and *z*[2 : 5] respectively, are shown in green. *z* has sufficient coverage because all five intervals include at least one of the starting positions of the reads.

### Assembly graphs

De Bruijn graphs, overlap graphs and string graphs are popular graph-based frameworks used for *de novo* genome assembly. Let *G*(*V*, *E*, *σ*, *w*) denote a directed assembly graph where vertices are labeled with strings, and edges indicate suffix-prefix overlap relations between the labels of connected vertices. Function *σ* : *V* → *Σ*^+^ assigns a string label to each vertex. Function *w* : *E* → ℕ assigns a weight to each edge. Weight of an edge *v*_1_ → *v*_2_ signifies the length of prefix of *σ*(*v*_1_) that is not in the suffix-prefix match. In a de Bruijn graph 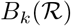, vertex set *V* corresponds to the set of all *k*-mer substrings in read set 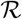, and an edge of weight one exists from vertex *v*_1_ to vertex *v*_2_ if and only if *σ*(*v*_1_)[2 : *k*] = *σ*(*v*_2_)[1 : *k* − 1]. This data structure is also sometimes referred to as node-centric de Bruijn graph [5].

Overlap graph 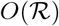 is a directed multi-graph where vertices correspond to sequence reads and edges correspond to suffix-prefix overlaps among the reads. A directed edge *v*_1_ → *v*_2_ with weight |*σ*(*v*_1_)|−*l* is drawn if and only if read *σ*(*v*_1_) has a suffix-prefix overlap of length *l* with read *σ*(*v*_2_). A subgraph of overlap graph 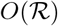 that contains only the edges representing suffix-prefix overlaps of length ≥ *k* is denoted as 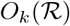. Myers [21] introduced a sparse variant of overlap graph structure called as string graph. Unlike overlap graphs which are defined as directed multi-graphs, a string graph, denoted as 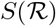 is a directed graph. A string graph includes only the longest suffix-prefix overlap between a read pair. Second, vertices associated with contained reads are excluded from a string graph. Finally, if vertex *v*_1_ connects to *v*_2_, *v*_2_ connects to *v*_3_ and *v*_1_ connects to *v*_3_ using edges *e*_1_, *e*_2_ and *e*_3_ respectively, and *w*(*e*_1_) + *w*(*e*_2_) = *w*(*e*_3_), then edge *e*_3_ is excluded from the string graph because it is transitively deducible. Removal of transitive edges is referred to as transitive sparsification. In practice, a string graph has significantly fewer edges compared to the overlap graph which enables computation of longer assembly contigs. A subgraph of string graph 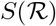 which only uses suffix-prefix overlaps of length ≥ *k* is denoted as 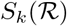.

### The circular strings spelled by graphs

By performing a closed walk in an assembly graph, one can “spell” a circular string. Suppose (*v*_1_, *e*_1_, *v*_2_, *e*_2_, …, *v*_*n*−1_, *e*_*n*−1_, *v*_*n*_), *n* > 1 is a sequence of vertices and edges in a closed walk such that edge *e*_*i*_ connects *v*_*i*_ to *v*_*i*+1_ and *v*_*n*_ = *v*_1_. The circular string spelled by this closed walk is 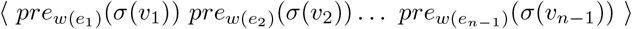. This is the circular string formed by concatenating labels of the vertices without spelling the overlapped substrings twice.

#### Definition 1.

*Given set of reads* 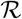, *coverage parameters* (*l*_1_, *l*_2_) *and chromosome length threshold ϕ, graph G*(*V, E, σ, w*) *is said to be **coverage-preserving** iff all candidate strings* ∈ 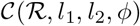 *can be spelled in the graph*.

## 3 Analysis of known graph models

### 3.1 De Bruijn graphs

#### Theorem 1.

*de Bruijn graph* 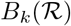 *is coverage-preserving for all k* ≤ *l*_2_ + 1.

*Proof*. If a circular string can be spelled in 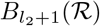, then it can certainly be spelled in 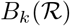, ∀*k* < *l*_2_ + 1. Therefore, it suffices to prove the claim for *k* = *l*_2_ + 1. We will prove that 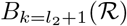 is coverage-preserving using contradiction. Suppose there exists a circular string 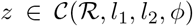 which cannot be spelled in 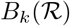. Let *k*_*i*_ (1 ≤ *i* ≤ *n*) denote the *k*-mer substring starting from position *i* in *z*. If all *k*_*i*_s exist in the set of all *k*-mer substrings extracted from read set 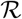, then it is trivial to construct a closed walk in 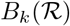 which spells *z*. Therefore, at least one of the *k*_*i*_s must be missing. Consider the minimum *i* for which this is true. Next, consider the (*l*_1_ – *l*_2_)-long interval ending at the position *i* in the circular string *z*. Based on the coverage assumption, there is at least one read ∈ 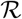 which matches a substring of length ≥ *l*_1_ of *z* starting in this interval. Such a read must contain *k*-mer *k*_*i*_ as its substring.

### 3.2 Overlap graphs

Unlike de bruijn graphs, overlap graphs require a more careful analysis due to variable-length suffix-prefix overlaps and variable-length vertex labels. The aim of this section is to prove the following.

#### Theorem 2.

*Overlap graph* 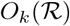 *is coverage-preserving for all k* ≤ *l*_2_.

If a circular string can be spelled in 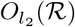, then it can be spelled in 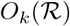, ∀*k* < *l*_2_. For a circular string 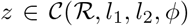, we propose an algorithm to identify a closed walk in 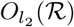 which spells *z*. For this, we first need some definitions. Let 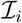 be the (*l*_1_ – *l*_2_)-long interval starting at position *i* in *z*, where *i* ranges from 1 to |*z*|. For each interval 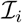, let 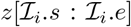 be the substring of length ≥ *l*_1_ which starts in interval 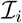 and matches a read 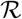. If there are multiple such substrings available, pick the ones with the least value of 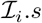, and then with the least value of 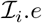. Let 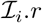 denote the read that matches substring 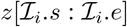. Recall that a circular string is equivalent to any of its cyclic rotated version, therefore, assume wlog that 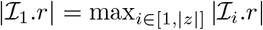. If 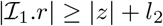, it is trivial to construct a valid closed walk by using a single vertex corresponding to read 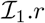. In the following, we will focus on the case when 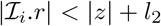 for all *i* ∈ [1, |*z*|]. The inequalities 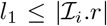 and 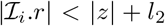 imply that the size of each interval, i.e., *l*_1_ – *l*_2_ is < |*z*|. Therefore, |*z*| > 1.

#### Lemma 1.

*For any two consecutive intervals* 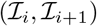 *in* 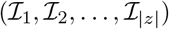,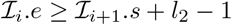.

*Proof*. All intervals including 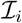 and 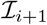 are of length *l*_1_ – *l*_2_. 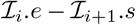 equals its minimum value *l*_2_ – 1 when (a) 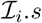 equals the first position in interval 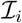, (b) 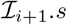 equals the last position in interval 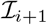, and (c) the substring 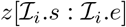 has minimum possible length *l*_1_.

Next we will identify a subsequence 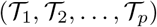 of 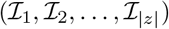 such that vertices corresponding to reads 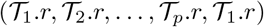 can be connected to form a valid closed walk which spells *z* (Figure 2). Let 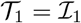. Suppose a partial subsequence 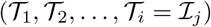 has been computed so far. Continue by selecting the first interval among 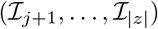 as 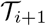 which satisfies the conditions 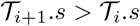 and 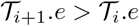. This selection procedure ensures that 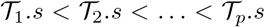 and 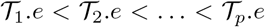. Suppose 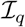 is the interval which ends at position 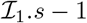 if 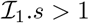 or else at position |*z*| if 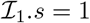.

**Fig. 2.**
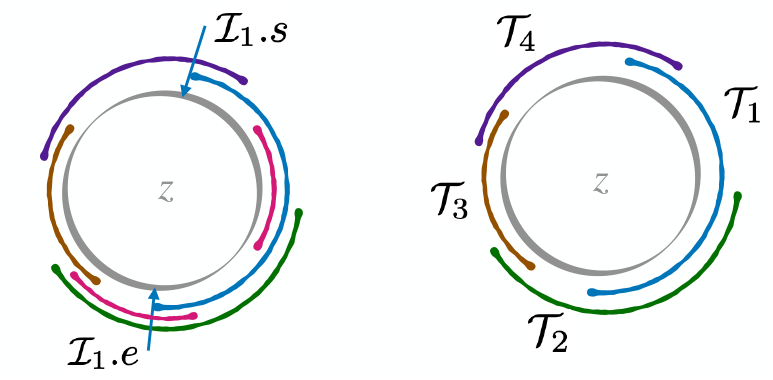
A graphic illustration of the subsequence computed from the complete sequence of intervals 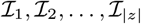. In the left figure, the gray-colored circle indicates circular string *z*, and the curved line segments indicate distinct substrings of *z* associated with intervals 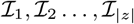. The right figure illustrates only the substrings associated with selected intervals in the subsequence 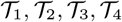.

#### Lemma 2.

*Length of the subsequence* 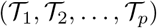 *identified by the above procedure is at least two*.

*Proof*. By contradiction. Assume *p* = 1, i.e., the computed subsequence is 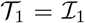. Based on our previous arguments, we have 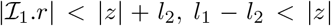 and |*z*| > 1. Interval 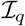 cannot contain position 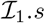 because size of each interval, i.e., *l*_1_ – *l*_2_, is < |*z*|. Therefore, 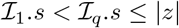. As interval 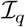 was not picked among the subsequence of intervals, 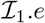 must be 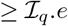. Next, we will identify the minimum possible value of 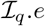 to get a lower bound of 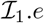. 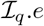 is lowest when the length of substring 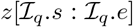 is *l*_1_ and 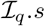 equals the first position in 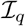. Using these conditions, observe that 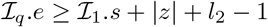, and therefore, 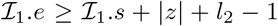. This implies that the length of substring 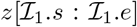 is ≥ |*z*| + *l*_2_. This is not possible when 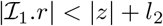.

#### Lemma 3.

*For any two consecutive intervals* 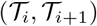 *in* 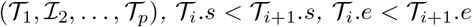 *and* 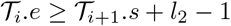.

*Proof*. The first two inequalities 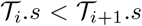 and 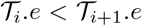 are guaranteed by the selection procedure of *T*_*i*+1_. Suppose 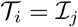 and 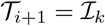. From Lemma 1, we know that 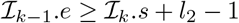. If *j* = *k* – 1, then inequality 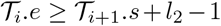 holds. Otherwise, *j* < *k* – 1 implies that 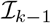 was not picked among the subsequence of intervals. This would happen in either of the following two situations (i) 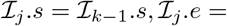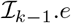, (ii) 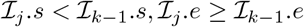. In either case, 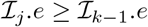. Therefore, 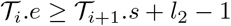.

#### Lemma 4.

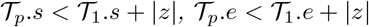 *and* 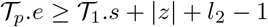.

*Proof*. The first inequality 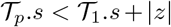 holds because 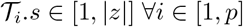. For the second inequality, recall that 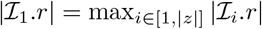. If 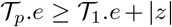, then 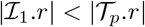 which cannot be true. The third inequality can be proved by using the arguments from the proof of Lemma 2. If *q* < |*z*|, then the intervals 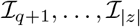 will contain 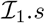 and won’t be selected in the subsequence. As a result, either *p* = *q* or *p* < *q*. In either case, 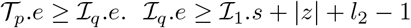 implies that 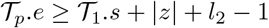.

Lemmas 3 and 4 collectively prove that vertices associated with the reads 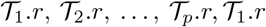 can be connected appropriately to build a valid closed walk in overlap graph 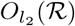. Along the walk, the non-overlapping prefix of read 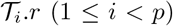 spells 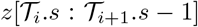. The prefix of the last read 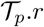 spells 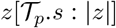. When 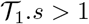, it also spells 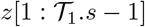. This proves Theorem 2.

### 3.3 String graphs

#### Theorem 3.

*String graph* 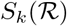 *is not guaranteed to be coverage-preserving for any value of k*.

*Proof*. A counter-example suffices to support the claim. Assume *l*_1_ = 6, *l*_2_ = 2 and *ϕ* = 10. Suppose count of reads 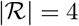, 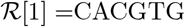, 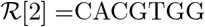, 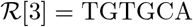 and 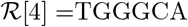 (Figure 3). Accordingly, the two candidate circular strings are ⟨CACGTGTG⟩ (covered by 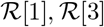) and ⟨CACGTGGG⟩ (covered by 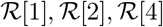). However, read 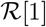 is contained in read 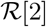 and the vertex corresponding to read 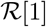 is excluded from the string graph. As a result, the first candidate ⟨CACGTGTG⟩ cannot be spelled.

**Fig. 3.**
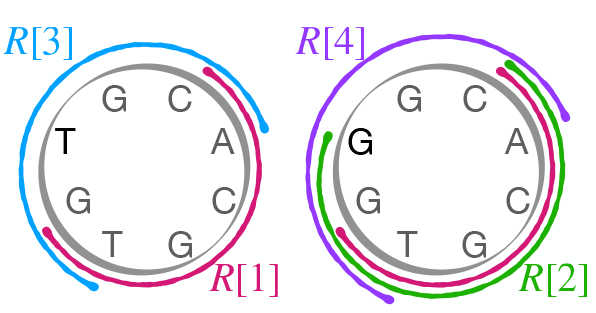
A visualization of the counter-example used to support Theorem 3.

The significance of the above result is that despite sufficient sequencing coverage, string graphs may lose coverage over the genome. This limitation has been observed in recent articles [7,22]. Accordingly, the following questions need to be addressed: (a) does the lack of the above guarantee affect quality of long-read assembly in practice? and (b) does there exist an alternate sparse graph model derived from an overlap graph whose size is similar to the string graph in practice, but guaranteed to be coverage-preserving?

## 4 An alternative framework for sparsification of overlap graphs

String graph is a sub-graph of an overlap graph that loses the coverage-preserving guarantee after pruning selected vertices and edges. In this section, we propose techniques to compute a directed multi-graph structure which is also a sub-graph of overlap graph, and it is guaranteed to be coverage-preserving. Specifically, we propose *safe* graph sparsification rules for vertex and edge removal from overlap graph 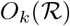, *k* ≤ *l*_2_ which guarantee that all circular strings ∈ 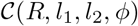 can be spelled in the sparse graph. Suppose 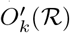 equals a subgraph of 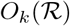 after performing a sequence of safe operations. Let *W*_*ϕ*_(*G*) denote the set of circular strings of length ≤ *ϕ* which can be spelled in graph *G*. From Theorem 2, we know that 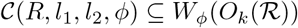. Therefore, the next two lemmas hold true.

### Lemma 5.

*Vertex u can be safely removed from graph* 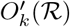 *if* 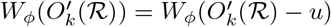.

### Lemma 6.

*Edge uv*_*l*_, *i.e., the edge from vertex u to vertex v of weight l can be safely removed from graph* 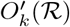 *if* 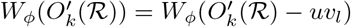.

### Lemma 7.

*If vertex v*_1_ *connects to v*_2_, *v*_2_ *connects to v*_3_ *and v*_1_ *connects to v*_3_ *using edges e*_1_, *e*_2_ *and e*_3_ *respectively such that w*(*e*_1_) + *w*(*e*_2_) = *w*(*e*_3_) *in graph* 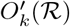, *then edge e*_3_ *can be safely removed*.

*Proof*. The above property follows from the transitive sparsification step during string graph construction [21]. A circular string spelled by a closed walk using edge *e*_3_ can be spelled by replacing *e*_3_ with edges *e*_1_ and *e*_2_ instead.

### Lemma 8.

*If a read* 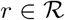 *is of length* < *l*_1_, *then the vertex corresponding to read r can be safely removed*.

*Proof*. The above property is implied from the proof of Theorem 2. Only the reads of length ≥ *l*_1_ were required in the algorithm to build a closed walk that spell each circular string ∈ 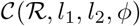.

One needs to be careful while filtering out contained reads as highlighted in the counter-example for Theorem 3. The next lemma suggests a condition for when it is safe to remove a contained read.

### Lemma 9.

*The vertex corresponding to a contained read* 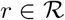 *can be safely removed from graph* 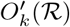 *if all the following conditions are satisfied*:

1. *Read r is a substring of exactly one candidate circular string* 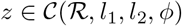.
2. *Read r matches a non-repetitive substring of z*.
3. *Graph* 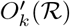 *contains a vertex corresponding to a parent read of r*.

*Proof*. Let *r*_*p*_ be a parent read of read *r* whose corresponding vertex exists in graph 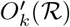. Denote the vertices corresponding to reads *r* and *r*_*p*_ as *u* and *u*_*p*_ respectively. Recall that all reads are substrings of the genome, and the genome is a subset of 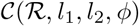. If *r* is a substring of exactly one circular string 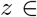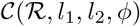, then *r*_*p*_ must be a substring of *z* and no other circular string in 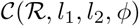. We need to show that *z* can still be spelled after removing vertex *u* from graph 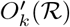. Suppose (*v*_1_, *e*_1_, *v*_2_, *e*_2_, …, *v*_*n*−1_, *e*_*n*−1_, *v*_*n*_), *n* > 1, *v*_*n*_ = *v*_1_ is a closed walk that includes vertex *u* to spell *z*. WLOG, assume that *v*_1_ = *u*. Note that neither of the vertices *v*_2_, *v*_3_, …, *v*_*n*−1_ equal *u* because read *r* matches a non-repetitive substring of *z*. Since *v*_1_ = *u*, the first character of *z* is spelled by using the first character of *r*, and *z*[1 : |*r*|] matches read *r*. Vertex *v*_1_ = *u* is said to span interval 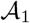 of length |*r*| starting from position one in *z*. Similarly, vertex *v*_2_ spans interval 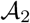 of length |*σ*(*v*_2_)| starting from position *w*(*e*_1_) + 1 in *z*. In general, each vertex *v*_*i*_ in the closed walk spans a unique interval 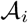 with starting position ∈ [1, |*z*|]. Next, we will construct a new closed walk to spell *z* that uses vertex *u*_*p*_ instead of vertex *u*. The interval spanned by vertex *u*_*p*_ in *z*, say 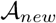, can be uniquely identified because substring *z*[1 : |*r*|] is non-repetitive. 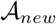 subsumes 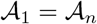 and possibly a few other adjacent intervals 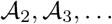 and 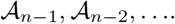. All vertices associated with the subsumed intervals can be removed and replaced with *u*_*p*_. With additional refinements, this procedure can be used to form a closed walk that spells *z* without using vertex *u*.

It would be ideal to construct a “minimal” sparse overlap graph using the above rules. However, implementing these rules in practice is not trivial. Suppose all sequencing errors are corrected by using an appropriate error-correction algorithm, we need to estimate *l*_1_, *l*_2_ and *ϕ* parameters. An efficient algorithm to compute set of candidate circular strings 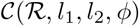 is unknown. Computing set of closed walks *W*_*ϕ*_(*G*) efficiently in large overlap graphs is also challenging.

## 5 A proof-of-concept implementation

We build a practical algorithm for overlap graph sparsification that removes transitive edges by using the property in Lemma 7, and filters out a subset of contained reads by using heuristics that are inspired from Lemmas 5, 9. Our implementation is designed for error-free long-read sequencing data sampled from both strands of DNA. Reverse complement of a string *x* is denoted as 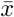. Reverse complements are handled by adding two separate vertices for each read: first for the read as given and second for its reverse complement. If vertex *v* corresponds to a read in either its forward or reverse-complemented orientation, then 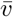 refers to the vertex corresponding to the read in the other orientation. Therefore, 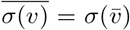. Graph *G*(*V, E, σ, w*) satisfies the following two properties: (i) 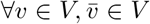, and (ii) For each edge *e* ∈ *E*, say from *v*_1_ to *v*_2_, an edge 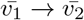 of weight |*σ*(*v*_2_) – (|*σ*(*v*_1_)| – *w*(*e*)) belongs to *E*.

Our implementation requires the following inputs from a user for the initial graph construction: (i) error-free long reads, (ii) minimum overlap length threshold, (iii) exact suffix-prefix overlaps, and (iv) exact match information of each contained read to each of its parent read. The minimum overlap length cutoff is set to 5 kbp by default. For our experiments, we used minimap2 [14] to compute all-vs-all read alignments. Overlaps available from minimap2 were filtered to satisfy the alignment length and 100% identity constraints. The first step in our implementation is to build an overlap graph from the input suffix-prefix overlaps, and separately label reads as either contained or non-contained. For each contained read *r*, a set of 4-tuples is used to save match information with the parent reads. For instance, tuple (*p, i, j*, +) indicates that read *r* matches substring *p*[*i* : *j*] of read 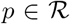. Similarly, tuple (*p, i, j*, ‒) indicates that read *r* matches substring 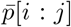, where 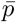 is the reverse complement of read 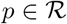.

### 5.1 Transitive sparsification of a directed multi-graph

The transitive sparsification property in Lemma 7 is inspired from the Myers’s string graph formulation [21]. However, their linear *O*(|*E*|) expected time algorithm [21] to label transitive edges is not applicable here because our graph is a multi-graph, i.e., graph can have multiple edges between the same source and target vertices. We use a simple *O*(|*E*|*D*^2^) time algorithm where *D* is the maximum out-degree of vertices in *G*.

**Algorithm 1:**
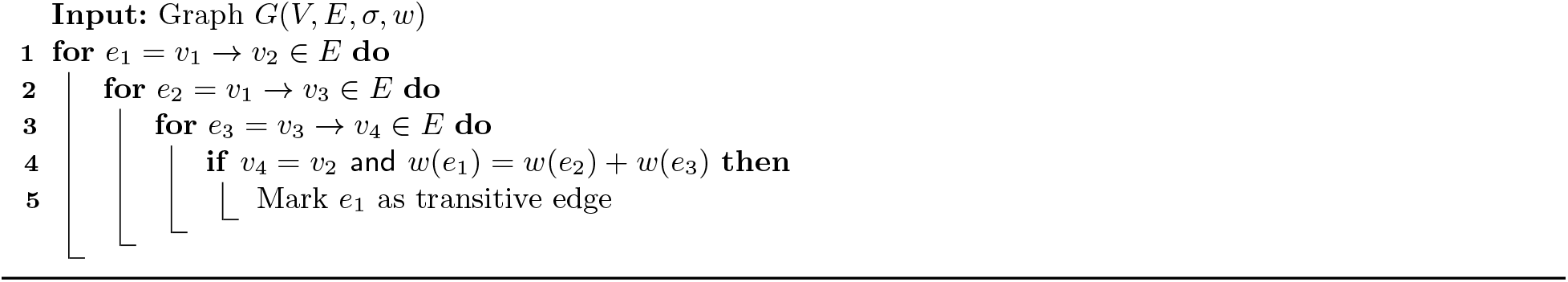
Labeling of transitive edges

#### 5.2 Remove non-repetitive haplotype-specific contained reads

Lemma 9 suggests that a contained read can be removed if (i) it is a substring of only a single haplotype, (ii) it is sampled from a non-repetitive genomic region, and (iii) at least one parent read is available in the graph. The third condition is automatically satisfied because there is at least one parent read of each read which is non-contained, and therefore, will not be removed. We propose a heuristic to check the first two conditions from all-versus-all read alignments computed using Hifiasm. For each read *r*, we inspect multiple sequence alignment (MSA) of read *r* and other reads overlapping with read *r*. We check if the count of reads in the MSA does not greatly exceed the sequencing coverage to ensure that the read is sampled from a non-repetitive genomic region. For a diploid genome, we additionally check for the presence of a heterozygous variant in the MSA (Supplementary Figure 2). If both the conditions are satisfied, we remove the two vertices corresponding to read *r* and its reverse complement respectively.

### 5.3 *k*-mer-based filtering heuristic for contained reads

We process the remaining contained reads using a second filter. In theory, we can try enumerating all possible string walks with and without contained read. Comparing the two will inform whether the read can be safely removed or not (Lemma 5). However, enumerating all possible string walks at chromosome length scale may not be computationally tractable in large graphs. We address this by implementing a *k*-mer-based heuristic. We first compute the set of *k*-mers observed in bounded-length string walks from the vertex associated with contained read (Supplementary Figure 1). The set is denoted as *κ*_1_. Next, we compute the union of set of *k*-mers observed in bounded-length string walks from the vertices associated with the parent reads of the contained read. This set is denoted as *κ*_2_. If *κ*_1_ ⊆ *κ*_2_, we mark the contained read for removal. This heuristic estimates whether there exists a string which cannot be spelled after removal of the contained read.

## 6 Experimental results

Our evaluation addresses two questions: (a) How many coverage gaps are introduced if we follow the standard string graph model, i.e., discard all contained reads during graph construction?, and (b) How well does the proposed implementation (hereon referred to as ContainX) perform when compared to string graph as well as other existing methods?

### 6.1 Experimental setup

#### Read simulation from human haploid and diploid genomes

We used two human genome assemblies for read simulation. The first is a CHM13 draft haploid genome assembly (v2.0) provided by the T2T consortium (GenBank id GCA 009914755.4) [22]. This haploid genome assembly of size 3.1 Gbp has 25 contigs and N50 length 50.6 Mbp. The second assembly is a long-read diploid genome assembly^1^ of HG002 human sample computed using Trio-hifiasm [3]. This diploid genome assembly of size 6.0 Gbp has 970 contigs and N50 length 57.8 Mbp. From these genome assemblies, we simulated eight *error-free* read sets whose length distributions are compatible with either real PacBio HiFi or ONT sequencing data. Seqrequester (https://github.com/marbl/seqrequester) is used to simulate reads from random start positions in both forward and reverse orientations. Seqrequester allows users to specify a desired read length distribution. We used HiFi and ONT read length distributions from publicly-available long-read datasets of HG02080 human sample [16]. Four long-read sets were simulated from haploid genome assembly with 20-fold coverage. The other four long-read sets were simulated from diploid genome assembly with 30-fold coverage (15-fold per haplotype). Length statistics and coverage information of the simulated read sets are shown in Table 1. The commands used to run the tools are listed in Supplementary Table 1.

**Table 1.**
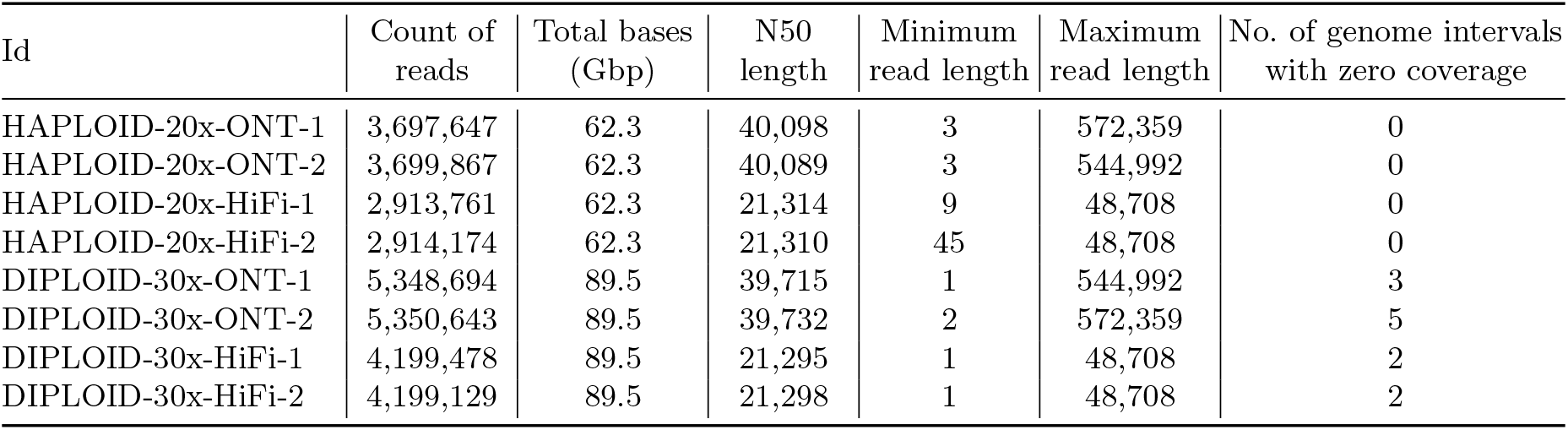
Properties of the simulated long-read sequencing datasets including their size and length statistics.

| Id | Count of reads | Total bases (Gbp) | N50 length | Minimum read length | Maximum read length | No. of genome intervals with zero coverage |
| --- | --- | --- | --- | --- | --- | --- |
| HAPLOID-20x-ONT-1 | 3,697,647 | 62.3 | 40,098 | 3 | 572,359 | 0 |
| HAPLOID-20x-ONT-2 | 3,699,867 | 62.3 | 40,089 | 3 | 544,992 | 0 |
| HAPLOID-20x-HiFi-1 | 2,913,761 | 62.3 | 21,314 | 9 | 48,708 | 0 |
| HAPLOID-20x-HiFi-2 | 2,914,174 | 62.3 | 21,310 | 45 | 48,708 | 0 |
| DIPLOID-30x-ONT-1 | 5,348,694 | 89.5 | 39,715 | 1 | 544,992 | 3 |
| DIPLOID-30x-ONT-2 | 5,350,643 | 89.5 | 39,732 | 2 | 572,359 | 5 |
| DIPLOID-30x-HiFi-1 | 4,199,478 | 89.5 | 21,295 | 1 | 48,708 | 2 |
| DIPLOID-30x-HiFi-2 | 4,199,129 | 89.5 | 21,298 | 1 | 48,708 | 2 |

### 6.2 Assessment of coverage gaps caused by removing contained reads

#### Benchmarking procedure

We developed a method to estimate coverage gaps introduced by removal of contained reads. We computed all-versus-all read overlaps by using minimap2 (v2.23) [14] to identify the set of contained reads. A read is labeled as contained if it matched a proper substring of a longer read. We can have false negatives because minimap2 uses *k*-mer-based heuristics. However, there cannot be false positives here because each overlap is verified using sequence alignment. In the second step, the set of non-contained reads was mapped to the genome. For the diploid genome, the set of non-contained reads was mapped to both paternal and maternal haplotypes, one by one. Minimap2 parameters were adjusted to enable reporting of multiple best alignments per read. Whenever a read aligned end-to-end against a genomic interval with 100% alignment identity, we recorded the interval. Finally, all the genomic intervals where no alignment coverage was observed were extracted using bedtools (v2.29.1). Not all of these can be considered as coverage gaps caused by removal of contained reads. The sequencing coverage and read lengths at the two extreme ends of each contig are expected to be lower than the whole-genome average due to edge effect. Therefore, the intervals that overlap with either the first 25 kbp or the last 25 kbp bases of a contig are not considered. Similarly, the intervals that overlap with segments of the genome with zero sequencing coverage are also not considered. The remaining intervals are the coverage gaps introduced by removal of contained reads.

#### Results

The count of coverage gaps observed in the four datasets are shown in Table 2. We observe that there are zero coverage gaps introduced in haploid genome for both ONT and HiFi read sets. In theory, haploid genomes can suffer coverage gaps due to identical repeats in the genome. However, removal of contained reads does not cause any harm in practice. In diploid genomes, however, we observe 1 and 46 − 54 coverage gaps introduced in HiFi and ONT datasets respectively. Compared to a haploid genome, coverage gaps in diploid genomes are more likely to happen because a longer read sampled from one haplotype can subsume all reads sampled from the homologous loci of the other haplotype, especially in regions with low heterozygosity. Moreover, ONT read length distribution is highly non-uniform which leads to a higher fraction of contained reads, and therefore, more coverage gaps. See Supplementary Figure 3 for long-read length distributions. We further investigated whether the observed gaps are clustered in a particular chromosome, but we do not observe such behavior. The coordinates of these coverage gaps in the HG002 assembly were mapped to the coordinates of GRCh38 human genome reference by using paftools [14]. The complete lists of these coordinates are provided in Supplementary Tables 2–5. We also evaluated heterozygosity rate by checking the count of heterozygous variants in these gaps. These gaps collectively span 1.26 million bases on GRCh38 genome reference when all the four diploid datasets are considered together. However, Dipcall [15] reported only 83 heterozygous variants which is an order of magnitude lower relative to the HG002 whole-genome average rate 0.1%. This observation confirms our expectation that the coverage gaps are more likely to happen in regions of low heterozygosity.

**Table 2.** Coverage gap statistics for the gaps introduced by removal of contained reads. Count of contained reads is estimated from all-to-all read alignments computed by using minimap2.

| Id | Count of contained reads | Coverage gap statistics |  |  |
| --- | --- | --- | --- | --- |
|  |  | Count | Mean length | Maximum length |
| HAPLOID-20x-ONT-1 | 3,160,926 | 0 | - | - |
| HAPLOID-20x-ONT-2 | 3,161,210 | 0 | - | - |
| HAPLOID-20x-HiFi-1 | 1,897,005 | 0 | - | - |
| HAPLOID-20x-HiFi-2 | 1,895,182 | 0 | - | - |
| DIPLOID-30x-ONT-1 | 4,550,993 | 46 | 11,227 | 53,490 |
| DIPLOID-30x-ONT-2 | 4,552,757 | 54 | 14,369 | 101,371 |
| DIPLOID-30x-HiFi-1 | 2,506,248 | 1 | 1,808 | 1,808 |
| DIPLOID-30x-HiFi-2 | 2,503,577 | 1 | 224 | 224 |

### 6.3 Evaluation of the proposed overlap graph sparsification algorithm

#### Benchmarking procedure

We evaluate the performance of our proof-of-concept implementation ContainX that determines a subset of contained reads to be retained. We tested ContainX using diploid datasets. We checked the performance by observing the following two parameters: (a) Count of coverage gaps caused by removing contained reads that are estimated to be redundant, and (b) count of “junction” reads, i.e., the reads which correspond to vertices having either > 1 incoming or > 1 outgoing edges in the graph. Retaining more contained reads implies that their corresponding vertices and edges are retained in the graph. This may result in more number of junction reads which can further result in shorter unitigs. It is important to look at the two parameters simultaneously because the first parameter can be optimized by removing all contained reads whereas the second parameter can be optimized by retaining all contained reads. An ideal solution should optimize both by doing a careful selection of contained reads. To measure the first parameter, we used minimap2 to compute end-to-end 100% identical read alignments of the retained contained reads against the two haplotypes. We checked if these alignments closed the coverage gaps that were previously caused by discarding all contained reads.

#### Competing methods

We compared ContainX with four other methods. The first method, called as *Retain-all*, simply retains all contained reads. The second method, called as *Remove-all*, discards all contained reads. The third method is from Hui *et al*. [9]. It removes a contained read *r* if two of its parent reads are *inconsistent*. Two parent reads are said to be inconsistent if, when aligned with respect to read *r*, they disagree at some base. We also included Hifiasm [3] in our benchmark. Unlike previous methods which stop at graph construction, Hifiasm is an full-fledged genome assembler that incorporates multiple heuristics (e.g., tip removal, bubble popping etc.) for graph sparsification. Hifiasm builds its initial overlap graph without using contained reads. It uses a heuristic to retain a few selected contained reads which can connect ends of two unitigs.

#### Results

We start with the evaluation of the first four methods; Hifiasm is discussed separately. The reads with no suffix-prefix overlap (above user-specified length) are excluded from the graph by default in all the four methods. In all four datasets, Retain-all method retained the highest number of contained reads as expected (Table 3). It resolved all coverage gaps, but the corresponding graph has significantly higher number of junction reads compared to Remove-all. Next, Hui-2016 filtering method appears to be conservative, i.e., it typically ends up retaining a majority of contained reads. In several cases, the contained read may have parent reads which agree at all bases (e.g., when they come from homozygous region of the genome). There may also be cases when there is only a single parent read available. In all such scenarios, Hui-2016 heuristic will choose to retain the contained reads. This heuristic resolved all the coverage gaps. Remove-all method has the fewest junction reads among the four methods but has the highest number of unresolved coverage gaps as expected. ContainX delivered favorable performance compared to the three methods. Using both datasets, ContainX retained 1 – 2% contained reads compared to Retain-all, yet it successfully resolved majority of the coverage gaps. ContainX resulted in only 2 – 5% junction reads when compared to Retain-all. This suggests that ContainX heuristics are promising in terms of improving assembly quality by avoiding coverage gaps. Finally, Hifiasm retained fewer contained reads than ContainX but it failed to resolve a majority of coverage gaps. This suggests that there is further scope to improve Hifiasm algorithm. The unitig graph of Hifiasm has the least number of junction reads because it does additional graph pruning which is necessary for computing longer unitigs. Incorporating ContainX heuristics inside Hifiasm code can be an interesting direction to explore in future.

**Table 3.**
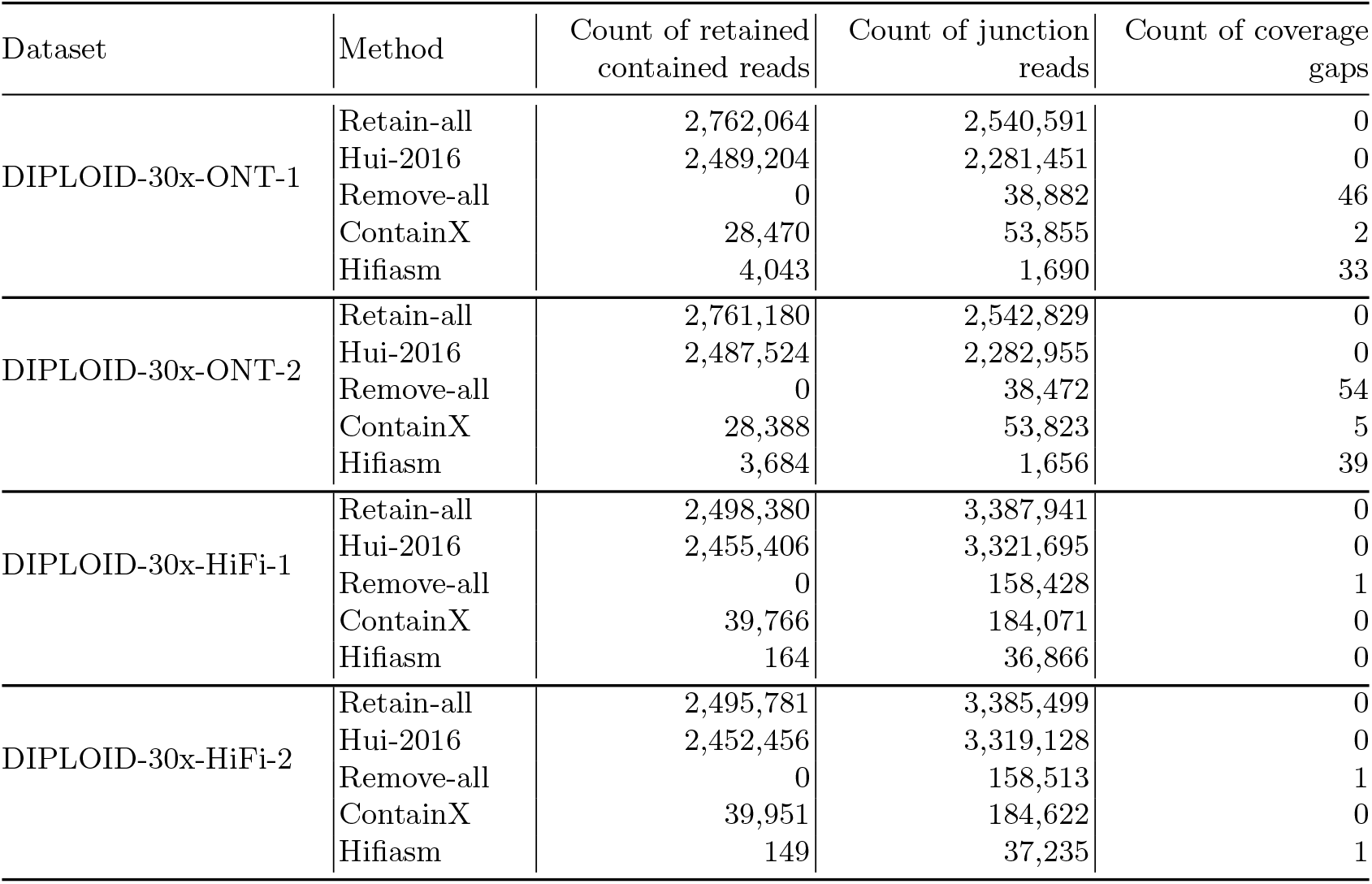
Performance comparison of ContainX with four other methods using long-read diploid datasets.

The proposed graph sparsification algorithms implemented in ContainX are easy-to-implement, fast and space-efficient. The *k*-mer-based heuristic uses multiple threads by considering each contained read independently. The time required by the ContainX heuristics using a multi-core AMD processor is about ten minutes. This is insignificant compared to the main time-consuming step during long-read assembly, i.e., computing all-vs-all read alignments.

## Acknowledgements

We thank Haowen Zhang, Mile Sikic and Robert Vaser for useful feedback. Haowen suggested the use of hifiasm code for one of the proposed heuristics. We thank Haoyu Cheng for helping us to modify hifiasm code. Brian Walenz helped to resolve an issue in Seqrequester code for long-read simulation. We also thank Sergey Nurk for highlighting this research problem during team discussions as part of the T2T consortium. This research is supported in part by funding from the Indian Institute of Science and allocation of U.S. National Energy Research Scientific Computing Center (NERSC) computing resources.

## Supplementary Information

**Fig. 1.**
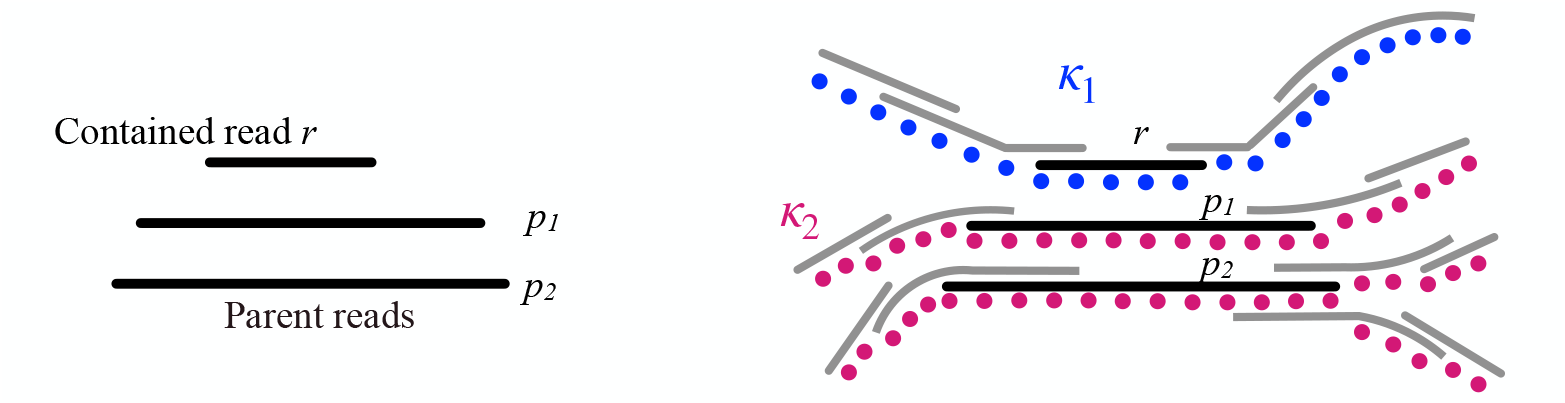
An illustration for how *k*-mer sets *κ*_1_ and *κ*_2_ are computed using bounded-length string walks from contained read *r* and its parent reads, respectively, in an overlap graph. Colored dots indicate the *k*-mers extracted from the strings.

**Fig. 2.**
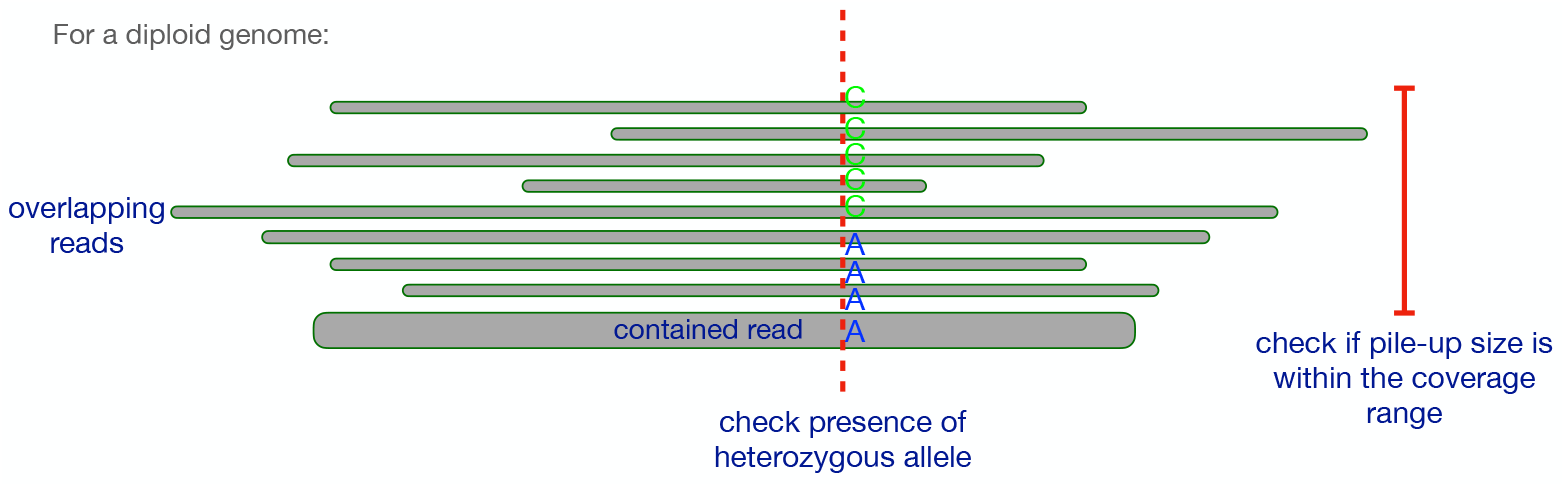
An illustration suggesting how multiple sequence alignment is used to detect non-repetitive haplotype-specific contained reads. In our implementation, the set of non-repetitive haplotype-specific reads is precomputed by using modified hifiasm code (https://github.com/cjain7/hifiasm/tree/hifiasm_dev_debug)

**Fig. 3.**
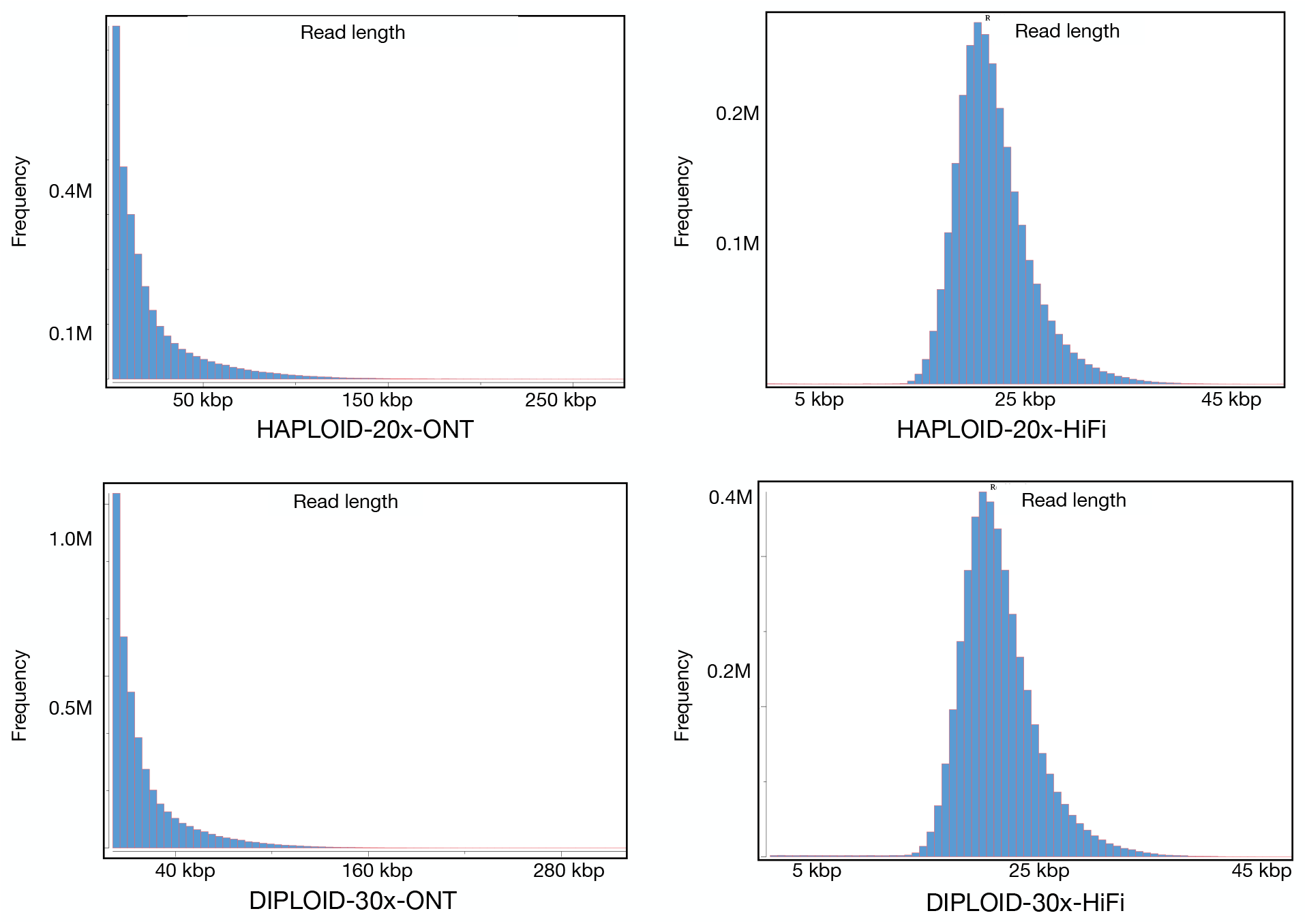
Histogram of the number of reads with each read length. Reads of length < 1000 bp are not considered. The above plots show that read length distribution of ONT datasets is highly skewed with a large number of reads of short length. HiFi read lengths are within a narrow range compared to ONT. These are known characteristics of the two sequencing technologies.

**Table 1.**
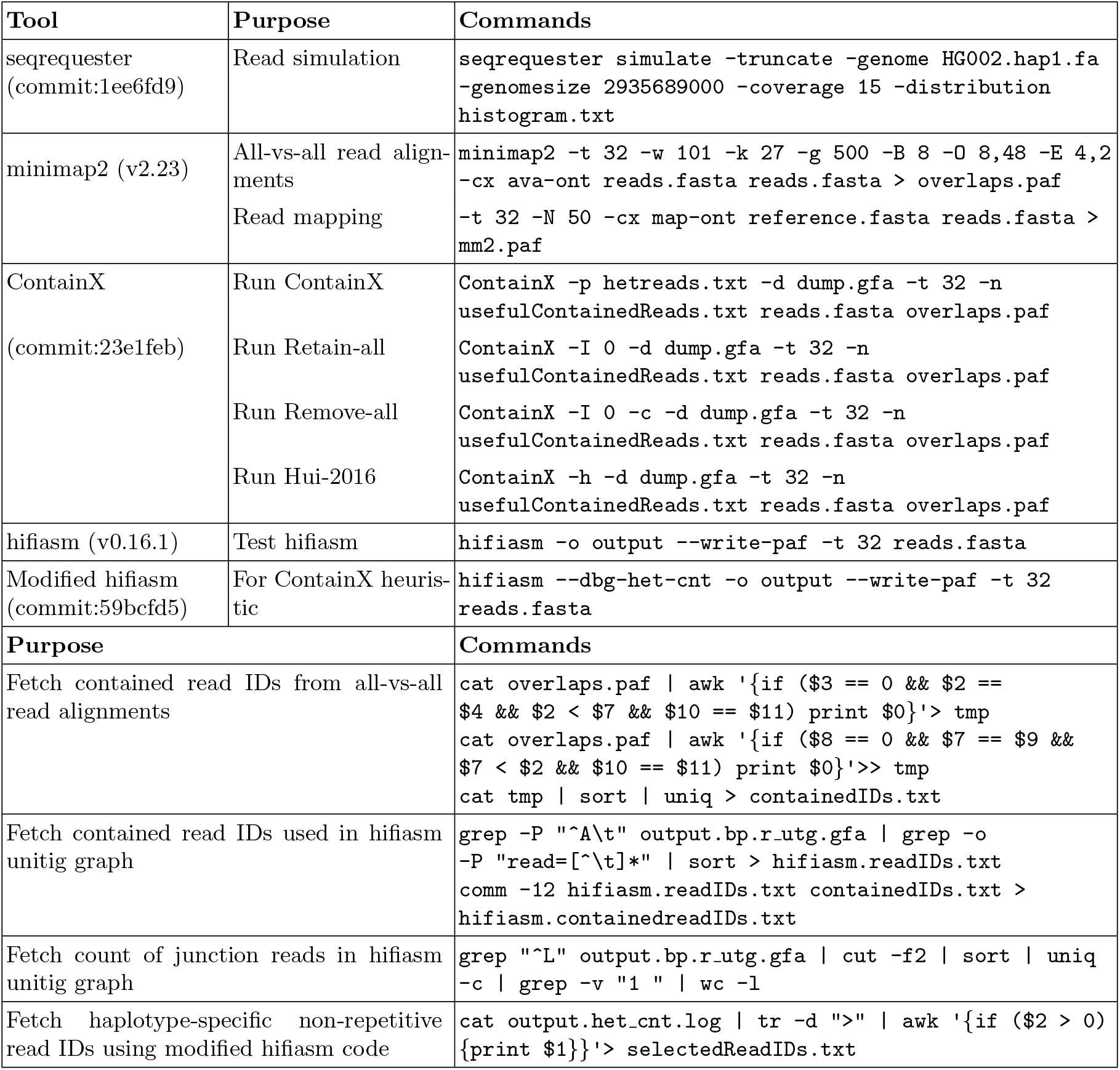
Some important commands and tools that were used in this study. Bash script files are also provided along with containX code.

**Table 2.**
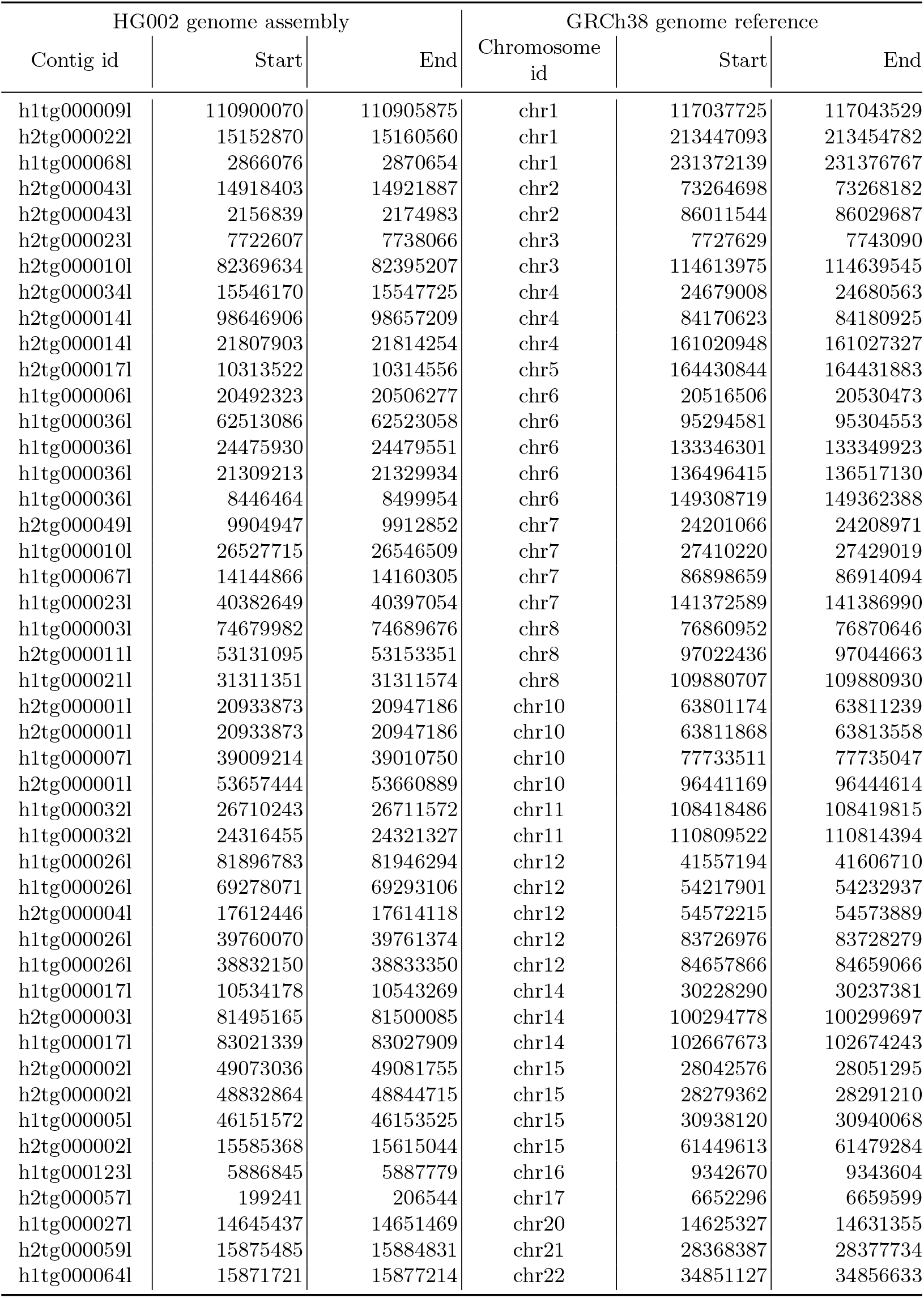
Start–stop coordinates of coverage gaps caused by removing contained reads in DIPLOID-30x-ONT-1 dataset. The coordinates in HG002 assembly are also translated (i.e., lifted over) to GRCh38 genome reference.

| HG002 genome assembly |  |  | GRCh38 genome reference |  |  |
| --- | --- | --- | --- | --- | --- |
| Contig id | Start | End | Chromosome id | Start | End |
| h1tg000009l | 110900070 | 110905875 | chr1 | 117037725 | 117043529 |
| h2tg000022l | 15152870 | 15160560 | chr1 | 213447093 | 213454782 |
| h1tg000068l | 2866076 | 2870654 | chr1 | 231372139 | 231376767 |
| h2tg000043l | 14918403 | 14921887 | chr2 | 73264698 | 73268182 |
| h2tg000043l | 2156839 | 2174983 | chr2 | 86011544 | 86029687 |
| h2tg000023l | 7722607 | 7738066 | chr3 | 7727629 | 7743090 |
| h2tg000010l | 82369634 | 82395207 | chr3 | 114613975 | 114639545 |
| h2tg000034l | 15546170 | 15547725 | chr4 | 24679008 | 24680563 |
| h2tg000014l | 98646906 | 98657209 | chr4 | 84170623 | 84180925 |
| h2tg000014l | 21807903 | 21814254 | chr4 | 161020948 | 161027327 |
| h2tg000017l | 10313522 | 10314556 | chr5 | 164430844 | 164431883 |
| h1tg000006l | 20492323 | 20506277 | chr6 | 20516506 | 20530473 |
| h1tg000036l | 62513086 | 62523058 | chr6 | 95294581 | 95304553 |
| h1tg000036l | 24475930 | 24479551 | chr6 | 133346301 | 133349923 |
| h1tg000036l | 21309213 | 21329934 | chr6 | 136496415 | 136517130 |
| h1tg000036l | 8446464 | 8499954 | chr6 | 149308719 | 149362388 |
| h2tg000049l | 9904947 | 9912852 | chr7 | 24201066 | 24208971 |
| h1tg000010l | 26527715 | 26546509 | chr7 | 27410220 | 27429019 |
| h1tg000067l | 14144866 | 14160305 | chr7 | 86898659 | 86914094 |
| h1tg000023l | 40382649 | 40397054 | chr7 | 141372589 | 141386990 |
| h1tg000003l | 74679982 | 74689676 | chr8 | 76860952 | 76870646 |
| h2tg000011l | 53131095 | 53153351 | chr8 | 97022436 | 97044663 |
| h1tg000021l | 31311351 | 31311574 | chr8 | 109880707 | 109880930 |
| h2tg000001l | 20933873 | 20947186 | chr10 | 63801174 | 63811239 |
| h2tg000001l | 20933873 | 20947186 | chr10 | 63811868 | 63813558 |
| h1tg000007l | 39009214 | 39010750 | chr10 | 77733511 | 77735047 |
| h2tg000001l | 53657444 | 53660889 | chr10 | 96441169 | 96444614 |
| h1tg000032l | 26710243 | 26711572 | chr11 | 108418486 | 108419815 |
| h1tg000032l | 24316455 | 24321327 | chr11 | 110809522 | 110814394 |
| h1tg000026l | 81896783 | 81946294 | chr12 | 41557194 | 41606710 |
| h1tg000026l | 69278071 | 69293106 | chr12 | 54217901 | 54232937 |
| h2tg000004l | 17612446 | 17614118 | chr12 | 54572215 | 54573889 |
| h1tg000026l | 39760070 | 39761374 | chr12 | 83726976 | 83728279 |
| h1tg000026l | 38832150 | 38833350 | chr12 | 84657866 | 84659066 |
| h1tg000017l | 10534178 | 10543269 | chr14 | 30228290 | 30237381 |
| h2tg000003l | 81495165 | 81500085 | chr14 | 100294778 | 100299697 |
| h1tg000017l | 83021339 | 83027909 | chr14 | 102667673 | 102674243 |
| h2tg000002l | 49073036 | 49081755 | chr15 | 28042576 | 28051295 |
| h2tg000002l | 48832864 | 48844715 | chr15 | 28279362 | 28291210 |
| h1tg000005l | 46151572 | 46153525 | chr15 | 30938120 | 30940068 |
| h2tg000002l | 15585368 | 15615044 | chr15 | 61449613 | 61479284 |
| h1tg000123l | 5886845 | 5887779 | chr16 | 9342670 | 9343604 |
| h2tg000057l | 199241 | 206544 | chr17 | 6652296 | 6659599 |
| h1tg000027l | 14645437 | 14651469 | chr20 | 14625327 | 14631355 |
| h2tg000059l | 15875485 | 15884831 | chr21 | 28368387 | 28377734 |
| h1tg000064l | 15871721 | 15877214 | chr22 | 34851127 | 34856633 |

**Table 3.** Start–stop coordinates of coverage gaps cause5d by removing contained reads in DIPLOID-30x-ONT-2 dataset.

| HG002 genome assembly |  |  | GRCh38 genome reference |  |  |
| --- | --- | --- | --- | --- | --- |
| Contig id | Start | End | Chromosome id | Start | End |
| h1tg000009l | 18493205 | 18504097 | chr1 | 24689133 | 24700023 |
| h1tg000009l | 86945009 | 86955138 | chr1 | 93087776 | 93097904 |
| h1tg000009l | 103508009 | 103514037 | chr1 | 109640847 | 109646875 |
| h1tg000012l | 64787435 | 64888806 | chr1 | 214650325 | 214751706 |
| h1tg000081l | 4756106 | 4770890 | chr2 | 73204290 | 73219075 |
| h1tg000081l | 11019519 | 11024843 | chr2 | 79484322 | 79489661 |
| h2tg000025l | 52097255 | 52097889 | chr2 | 212663821 | 212664455 |
| h1tg000001l | 92817961 | 92818455 | chr2 | 223349132 | 223349626 |
| h1tg000004l | 8873984 | 8877229 | chr3 | 8874669 | 8877914 |
| h2tg000184l | 232777 | 248699 | chr3 | 74871732 | 74887654 |
| h2tg000034l | 4797508 | 4823424 | chr4 | 13932709 | 13958691 |
| h1tg000042l | 114230605 | 114234253 | chr4 | 73904437 | 73908085 |
| h1tg000042l | 104113807 | 104122161 | chr4 | 84022694 | 84031047 |
| h2tg000014l | 92028013 | 92033684 | chr4 | 90811125 | 90816796 |
| h2tg000014l | 82459823 | 82460036 | chr4 | 100390877 | 100391090 |
| h2tg000014l | 70702729 | 70707859 | chr4 | 112148705 | 112153851 |
| h1tg000042l | 15091152 | 15118100 | chr4 | 173050753 | 173077668 |
| h1tg000016l | 91734766 | 91742004 | chr5 | 83043812 | 83051050 |
| h2tg000017l | 90073347 | 90073761 | chr5 | 84683858 | 84684272 |
| h1tg000006l | 20609242 | 20654818 | chr6 | 20633454 | 20679068 |
| h2tg000018l | 25740048 | 25747436 | chr6 | 33724209 | 33731597 |
| h1tg000006l | 50115047 | 50118074 | chr6 | 50067746 | 50070778 |
| h2tg000021l | 5153975 | 5157462 | chr6 | 64965355 | 64968838 |
| h1tg000036l | 90457178 | 90459095 | chr6 | 67337810 | 67339727 |
| h1tg000010l | 20186319 | 20234628 | chr7 | 21071775 | 21120082 |
| h2tg000049l | 12785813 | 12787547 | chr7 | 21326099 | 21327801 |
| h2tg000049l | 9850119 | 9868340 | chr7 | 24245575 | 24263817 |
| h1tg000145l | 1110399 | 1110479 | chr7 | 67358338 | 67358418 |
| h2tg000006l | 26473188 | 26544931 | chr7 | 132862940 | 132934640 |
| h2tg000006l | 26231773 | 26253441 | chr7 | 133155335 | 133177002 |
| h1tg000023l | 40495078 | 40505896 | chr7 | 141485319 | 141496136 |
| h1tg000003l | 46208131 | 46209187 | chr8 | 48451474 | 48452530 |
| h1tg000003l | 64446587 | 64457392 | chr8 | 66634485 | 66645289 |
| h2tg000011l | 22611658 | 22627754 | chr8 | 67277359 | 67293455 |
| h1tg000059l | 13139707 | 13143077 | chr9 | 28243394 | 28246761 |
| h2tg000041l | 37246637 | 37249901 | chr9 | 37218351 | 37221615 |
| h1tg000007l | 11801161 | 11811351 | chr10 | 104918759 | 104928949 |
| h2tg000038l | 50521189 | 50532240 | chr11 | 39142190 | 39153262 |
| h1tg000011l | 44597468 | 44597862 | chr11 | 46713582 | 46713976 |
| h2tg000007l | 24589917 | 24627125 | chr11 | 110494690 | 110531902 |
| h2tg000037l | 1789315 | 1816591 | chr13 | 19277044 | 19304307 |
| h2tg000037l | 12969569 | 12969897 | chr13 | 30439954 | 30440282 |
| h2tg000037l | 19506191 | 19509196 | chr13 | 36983851 | 36986858 |
| h1tg000022l | 42499978 | 42505754 | chr13 | 44248785 | 44254562 |
| h2tg000037l | 30583828 | 30599471 | chr13 | 48052070 | 48067716 |
| h2tg000037l | 33124120 | 33126064 | chr13 | 50591608 | 50593552 |
| h1tg000033l | 20360199 | 20363658 | chr13 | 107116486 | 107119945 |
| h2tg000003l | 26788478 | 26790995 | chr14 | 45599428 | 45601940 |
| h2tg000002l | 2549333 | 2575965 | chr15 | 74425030 | 74451662 |
| h1tg000005l | 1570767 | 1570875 | chr15 | 75451659 | 75451767 |
| h1tg000069l | 22949634 | 22972649 | chr16 | 78611705 | 78634697 |
| h1tg000030l | 14192878 | 14193075 | chr17 | 3288793 | 3288990 |
| h2tg000224l | 30443 | 72254 | chr19 | 43484700 | 43525330 |
| h1tg000060l | 29858829 | 29903277 | chr21 | 14516790 | 14561239 |

**Table 4.** Start–stop coordinates of coverage gaps caused by removing contained reads in DIPLOID-30x-HiFi-1 dataset.

| HG002 genome assembly |  |  | GRCh38 genome reference |  |  |
| --- | --- | --- | --- | --- | --- |
| Contig id | Start | End | Chromosome id | Start | End |
| h1tg000036l | 57223158 | 57224966 | chr6 | 100588432 | 100590240 |

**Table 5.** Start–stop coordinates of coverage gaps caused by removing contained reads in DIPLOID-30x-HiFi-2 dataset.

| HG002 genome assembly |  |  | GRCh38 genome reference |  |  |
| --- | --- | --- | --- | --- | --- |
| Contig id | Start | End | Chromosome id | Start | End |
| h2tg000002l | 32863923 | 32864147 | chr15 | 44197800 | 44198024 |

## Footnotes

1 ftp://ftp.dfci.harvard.edu/pub/hli/hifiasm-phase/v2/HG002.hifiasm.trio.0.16.1.hap1.fa.gz ftp://ftp.dfci.harvard.edu/pub/hli/hifiasm-phase/v2/HG002.hifiasm.trio.0.16.1.hap2.fa.gz

